# From Lipid Dynamics to Precision Predictions: A New Approach Methodology for Precision Modeling of Phosphoinositide Signaling

**DOI:** 10.1101/2025.07.03.663046

**Authors:** Gonzalo Hernandez-Hernandez, Mindy Tieu, Pei-Chi Yang, Oscar Vivas, Timothy J. Lewis, L. Fernando Santana, Colleen E. Clancy

## Abstract

Precision medicine requires models that can translate rich molecular measurements into individualized predictions of biological response. This challenge is particularly acute for phosphoinositide signaling disorders that often exhibit cell-type-specific responses to identical genetic or pharmacological perturbations. Here, we develop a New Approach Methodology (NAM) demonstrating that basal phosphoinositide pool composition, determined by the size of the PI(4)P reserve, determines the robustness of lipid signaling. The NAM comprises a core kinetic model of phosphatidylinositol (PI), phosphatidylinositol 4-phosphate (PI(4)P), phosphatidylinositol 4,5-bisphosphate (PI(4,5)P_2_), and inositol 1,4,5-trisphosphate (IP_3_) dynamics. The model also incorporates phospholipase C (PLC)-mediated hydrolysis and phosphatase-mediated turnover and explicitly accounts for IP_3_ biosensor binding during parameter optimization. Parameters were optimized using experimental measurements from superior cervical ganglion (SCG) neurons and validated against independent dose-dependent PI(4,5)P_2_ depletion data. Local and global sensitivity analyses were performed to identify the dominant parameter drivers of pathway behavior. These sensitivity relationships were then used to generate a population of model variants that captured phosphoinositide dynamics observed in tsA201 cells, human neuroblastoma cells, and hippocampal neurons. To infer cell-specific models, we developed two complementary inverse methods: sensitivity fingerprinting derived from mechanistic model sensitivities and a neural network trained on synthetic phosphoinositide time series. Both approaches reproduced experimental PI(4)P, PI(4,5)P_2_, and IP3 dynamics across cell types while preserving the baseline model structure. Importantly, the inferred models predicted experimentally observed differential vulnerability to kinase perturbation without additional fitting. Hippocampal neurons with large basal pools of PI(4)P maintained PI(4,5)P_2_ and IP_3_ signaling under phosphatidylinositol 4-kinase alpha (PI4KA) inhibition, whereas cells with small basal PI(4)P pools exhibited signaling failure. Simulations of PI4KA and phosphatidylinositol-4-phosphate 5-kinase type 1 gamma (PIP5K1C) loss-of-function mutations under repeated stimulation further revealed progressive signaling collapse in small-pool neurons but sustained function in large-pool neurons, demonstrating that basal lipid composition can determine genetic vulnerability. Together, this NAM provides a predictive, cell-specific framework for translating dynamic lipid measurements into mechanistic models that support precision medicine applications in phosphoinositide-related disorders.

## Introduction

Precision medicine aims to tailor treatment by accounting for patient-to-patient variation in genes, proteins, and cellular physiology, differences that are averaged in traditional approaches. Achieving this goal requires not only richer molecular measurements but also dynamic, mechanistic models that can translate measurements into individualized predictions of biological response. Computational New Approach Methodologies (NAMs) provide a path toward this translation by replacing animal experiments with predictive, data-driven modeling frameworks that can ultimately be applied to patient and cell-specific inference. This need is acute in lipid signaling disorders, where nonlinear pathway dynamics and chemically diverse lipid intermediates make mechanistic interpretation difficult without explicit dynamic models (Zandl-Lang et al., 2023). Accordingly, lipid signaling represents both a major challenge and a major opportunity for precision medicine.

Lipidomics represents a critical frontier for precision medicine. Phosphatidylinositol 4,5 bisphosphate (PI(4,5)P_2_) is fundamental to this effort because it acts as a central regulator of cellular signaling, membrane dynamics, and metabolism, and these processes can vary substantially across individuals (Balla, 2013; Viaud et al., 2016). Lipid dysregulation contributes to diseases including cancer, neurodegeneration, and metabolic disorders, making lipid pathways valuable both as biomarkers and as therapeutic targets (Burke et al., 2023). However, lipids remain challenging to study in practice because of their structural diversity, transient dynamics, and strong context dependence, requiring sophisticated tools and multidisciplinary strategies.

PI(4,5)P_2_ plays a central role in plasma membrane signaling through a tightly coupled and highly dynamic metabolic network. PI(4,5)P_2_ synthesis proceeds through sequential phosphorylation of phosphatidylinositol (PI) to phosphatidylinositol-4-phosphate (PI(4)P) and then to PI(4,5)P_2_, catalyzed by PI 4-kinase (PI4K) and PIP 5-kinase (PIP5K), respectively, with each step consuming ATP. PI is the most abundant phosphoinositide and serves as the precursor for the entire pathway; the PI pool is maintained through resynthesis in the endoplasmic reticulum and transport to the plasma membrane via lipid transfer proteins at membrane contact sites (Grabon et al., 2017; Kim et al., 2015; Pemberton et al., 2020). PI(4)P functions as an obligate intermediate whose concentration is dynamically coupled to that of PI(4,5)P_2_, such that changes in flux through PI(4)P rapidly propagate to PI(4,5)P_2_. During receptor-mediated activation of phospholipase C (PLC), PI(4)P and PI(4,5)P_2_ are depleted in parallel on a seconds timescale, reflecting the continuous consumption of PI(4)P to sustain PI(4,5)P_2_ synthesis while PI(4,5)P_2_ is simultaneously hydrolyzed by PLC (Falkenburger et al., 2010; Willars et al., 1998; Xu et al., 2003). Hydrolysis of PI(4,5)P_2_ generates diacylglycerol (DAG) and inositol trisphosphate (IP3), two second messengers with distinct temporal and spatial dynamics: IP_3_ diffuses rapidly through the cytoplasm to trigger calcium release from intracellular stores, whereas DAG remains membrane-associated and activates protein kinase C), two second messengers that propagate downstream responses (Berridge, 1993, 2009; Berridge & Irvine, 1984, 1989). Because IP_3_ has a short half-life (approximately 5-30 seconds) (Falkenburger et al., 2010), its concentration tracks the instantaneous rate of PLC activity rather than cumulative production, making it a direct readout of pathway flux. Although PI(4,5)P_2_ plays a structural role in maintaining the compositional and functional organization of the inner leaflet of the plasma membrane and supporting membrane trafficking, ion channel function, and cytoskeletal organization (Hille et al., 2015; Dickson and Hille, 2019), the pathway is fundamentally governed by rapid turnover and tightly regulated feedback. The phosphoinositide pathway is inherently nonlinear because its intermediates are linked through finite substrate pools and competing enzymatic reactions. Consumption of PI(4,5)P_2_ during PLC activation simultaneously reduces membrane lipid abundance, alters substrate availability for downstream reactions, and changes the rate at which signaling molecules are produced. As a result, identical changes in kinase activity can generate markedly different signaling outcomes depending on the starting lipid composition of the cell. These nonlinear dynamics make the outcomes of genetic or pharmacological perturbations difficult to predict without explicit quantitative models.

Here, we utilized measurements of PI(4)P, PI(4,5)P_2_, and IP_3_ with sufficient temporal resolution for kinetic parameter extraction across multiple cell systems: superior cervical ganglion (SCG) neurons (Kruse et al., 2016), tsA201 cells (Dickson et al., 2013), human neuroblastoma cells (Willars et al., 1998), and hippocampal neurons (de la Cruz et al., 2022). Despite differences in cell type, receptor systems, and experimental protocols, the datasets revealed a qualitatively consistent pathway architecture: rapid PI(4)P and PI(4,5)P_2_ depletion, coordinated IP_3_ production, and gradual recovery, all following the same sequence of biochemical reactions. However, the magnitude and kinetics of these responses varied substantially across cell types, reflecting quantitative differences in enzyme expression, lipid pool sizes, and cellular geometry. The shared pathway topology suggested that a common biochemical architecture governs phosphoinositide metabolism, with quantitative differences arising from systematic parameter variation rather than distinct pathway organization. The observation motivated a strategy: optimize a kinetic model to one well-characterized reference dataset, then use sensitivity analysis to guide parameter variation across other cell types.

We next developed a NAM that transforms optimized kinetic models into cell-specific inference tools through systematic sensitivity analysis. The framework integrates ordinary differential equations with multi-start Nelder-Mead optimization (Hernandez-Hernandez et al., 2024; Kernik et al., 2019; Moreno et al., 2016) to establish a reference baseline, then employs local and global Partial Rank Correlation Coefficient (PRCC) sensitivity analysis to map how individual kinetic parameters control distinct signaling features. The sensitivity relationships define a low-dimensional parameter structure that supports three complementary analyses: population modeling, sensitivity-guided inference of cell-specific kinetic parameters, and neural network-based parameter estimation from phosphoinositide time-series data. This integration of mechanistic modeling and machine learning provides an interpretable and scalable framework for constructing cell-specific phosphoinositide models.

We applied the framework to phosphoinositide signaling by optimizing a kinetic model for SCG neurons, then extending it across cell types. The analysis revealed that basal lipid pool composition determines signaling robustness and disease vulnerability. Simulations of loss-of-function mutations in PI4KA and PIP5K1C, lipid kinases implicated in neurodevelopmental and neuromuscular disorders (Burke et al., 2023; Narkis et al., 2007; Salter et al., 2021; Verdura et al., 2021), showed that identical genetic perturbations produce severe signaling impact in cells with small lipid reserves but allow partial compensation in cells with large reserves. The cell type-dependent predictions demonstrate how computational NAMs can translate genetic variants into mechanistic predictions by linking genotype to cellular phenotype. While demonstrated here with established cell lines, the NAM could be extended to enable patient-specific kinetic fingerprinting: measuring phosphoinositide dynamics in patient-derived cells, inferring cell-specific parameters via sensitivity-guided analysis, and predicting vulnerability to genetic or pharmacological perturbations.

## Results

We developed a New Approach Methodology (NAM) to transform an optimized kinetic signaling model into a mechanistic inference framework, enabling systematic analysis of cell-specific variability. Conceptually, the framework addresses four questions: What is the baseline signaling behavior? Which parameters control observable features? How do those parameters differ between cell types? Do those inferred differences predict responses to perturbation? The overall strategy follows four steps: (1) optimize a baseline kinetic model to a reference cell type; (2) use sensitivity analysis to map which parameters control which observable features; (3) use that mapping to infer cell-type-specific models from experimental time series, either manually or via a neural network; and (4) test whether the inferred models correctly predict the effects of genetic perturbations. We demonstrate this strategy using phosphoinositide signaling to generate cell-specific models relevant to precision medicine applications.

We constructed a kinetic model of PI(4)P, PI(4,5)P_2_, and IP_3_ dynamics using differential equations governed by six enzymatic rate constants constrained by biochemical conservation. The model represents lipid phosphorylation, dephosphorylation, PLC-mediated hydrolysis, and IP_3_ degradation, while explicitly accounting for IP_3_ biosensor binding **(Figure 1)**. IP_3_ is measured experimentally with the LIBRAvIII biosensor, which is a genetically encoded fluorescent reporter that binds free IP_3_ and sequesters a fraction it. The model accounts for the sequestration by converting free IP_3_ to sensor-bound IP_3_, to allow model comparison with experimental fluorescence signals **(Figure 1B)**. Unless otherwise noted, IP_3_ traces throughout represent biosensor-bound IP_3_ (IP_3_-LIBRAvIII), denoted as IP_3_*, which is the experimentally measured quantity in most datasets. PI(4)P and PI(4,5)P_2_ model outputs are compared directly with experimentally measured lipid reporter signals across all cell types.

**Figure 1.**
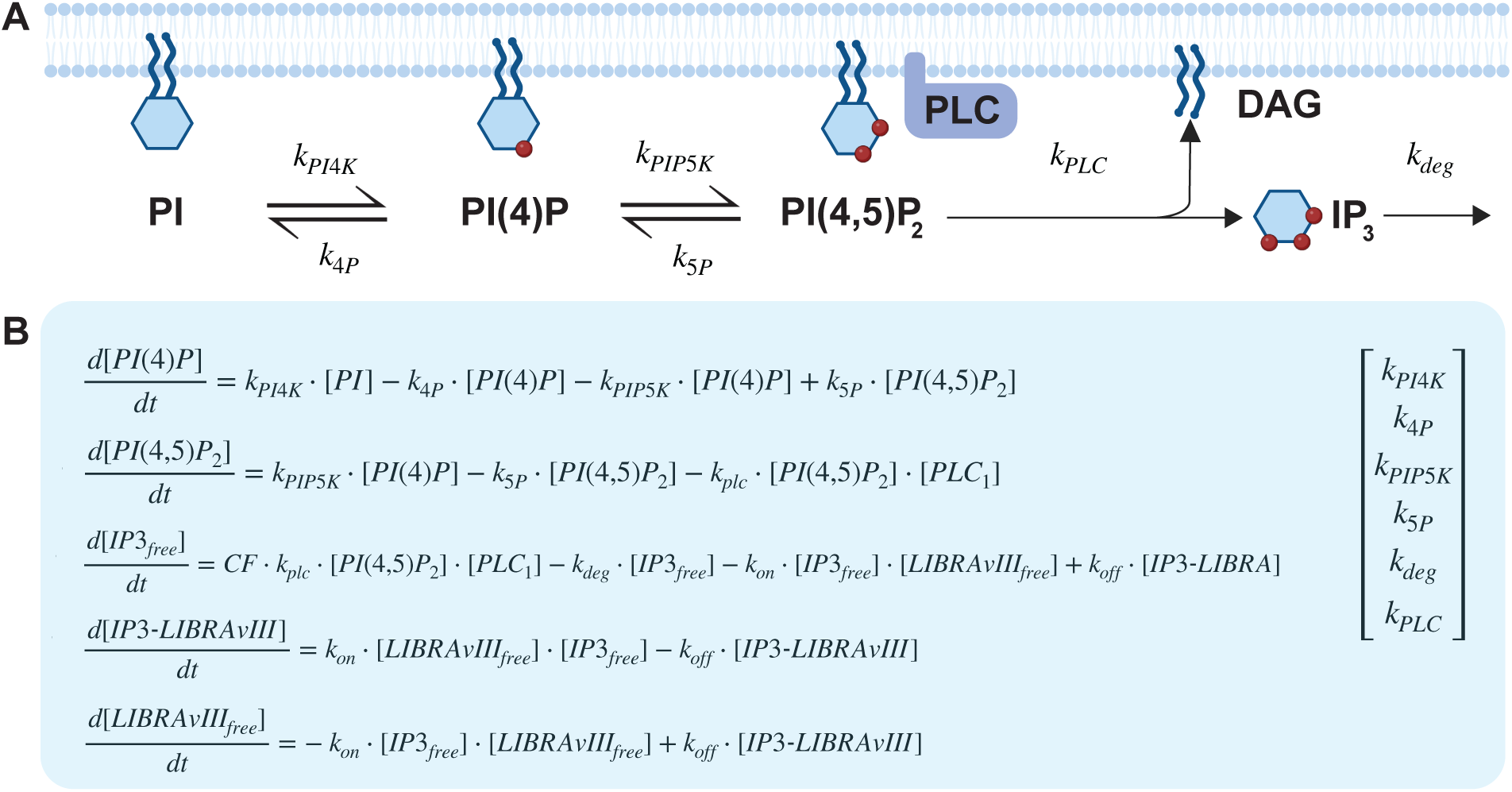
Phosphoinositide signaling pathway and kinetic model. **A**) Phosphoinositide signaling pathway schematic. Phosphatidylinositol (PI) undergoes sequential phosphorylation to phosphatidylinositol 4-phosphate (PI(4)P) and phosphatidylinositol 4,5-bisphosphate (PI(4,5)P_2_), with dephosphorylation mediated by 4-phosphatase and 5-phosphatase activity. PLC hydrolyzes PI(4,5)P_2_ to generate membrane-bound DAG and soluble inositol 1,4,5-trisphosphate (IP_3_). (**B**) Differential equations describing species concentrations (left) with dynamics governed by six kinetic rate constants: k_PI4K_, k_4P_, k_PIP5K_, k_5P_, k_deg_, k_PLC_. The model explicitly includes LIBRAvIII biosensor binding to IP_3_, which is accounted for during parameter optimization.

The model was optimized using experimental time-course data from superior cervical ganglion (SCG) neurons (**Figure 2A**), where 10 μM Oxo-M applied for 20s activates muscarinic M1 receptors and drives PLC-mediated hydrolysis of PI(4,5)P_2_, producing coordinated depletion of PI(4)P and PI(4,5)P_2_ and generation of IP_3_. Six kinetic rate constants (k_PI4K_, k_4P_, k_PIP5K_, k_5P_, k_PLC_, and k_deg_) were estimated using the Nelder-Mead optimization algorithm (**Table 1**). The optimized model reproduced the timing and magnitude of PI(4)P (**Figure 2A-i**), PI(4,5)P_2_ (**Figure 2A-ii**), and IP_3_ (**Figure 2A-iii**), transients observed experimentally. The model reproduced experimental dynamics with normalized root mean square errors of 8.9% for PI(4)P, 3.2% for PI(4,5)P_2_, and 4.5% for IP_3_, well below the 10% threshold typically considered acceptable for biological time-series fitting.

**Figure 2.**
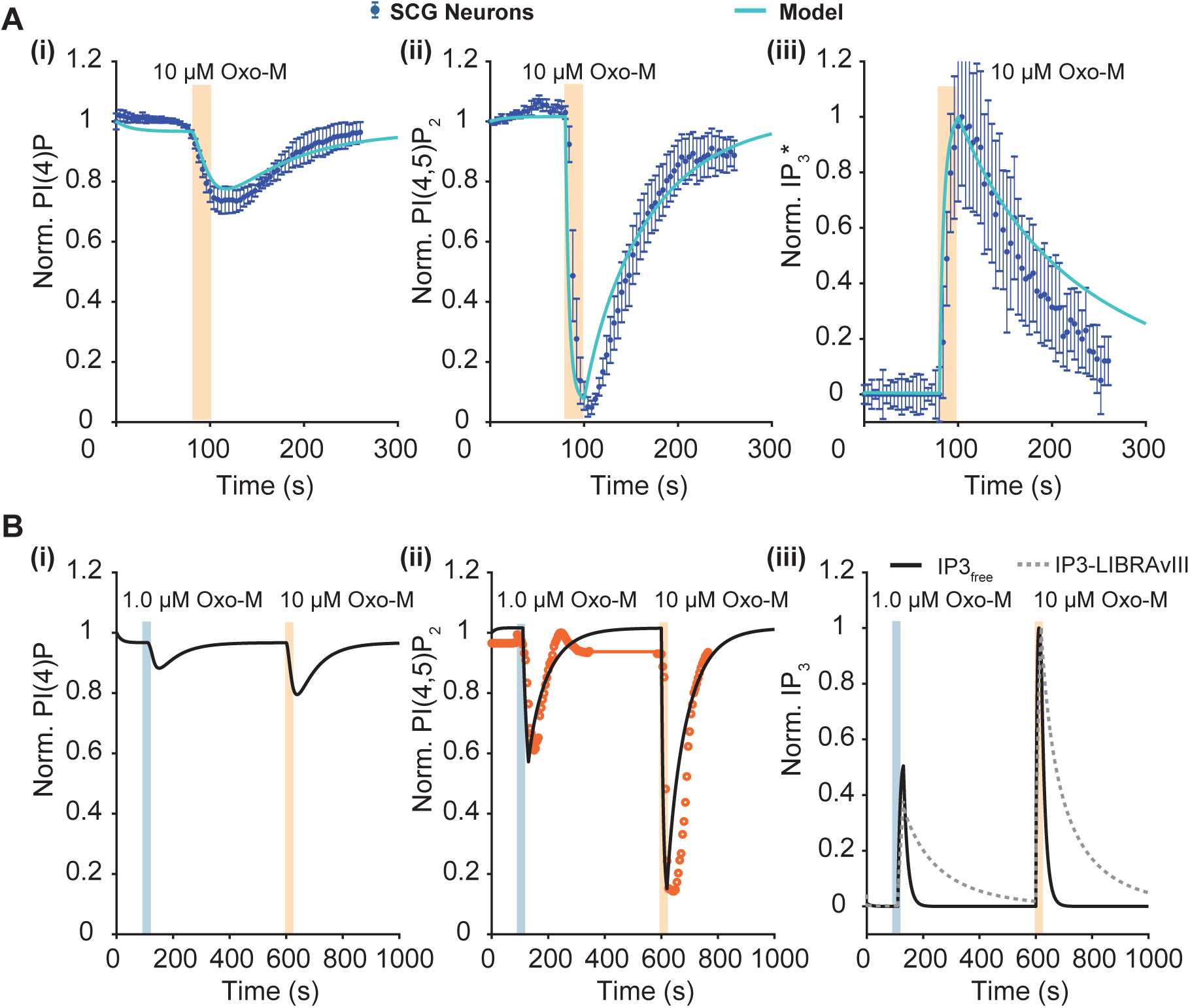
Model optimization and validation in superior cervical ganglion (SCG) neurons. (**A**) Parameter optimization from experimental data (blue symbols; Kruse et al., 2016) following 10 μM Oxo-M for 20 s (pink shading). Six kinetic rate constants (k_PI4K_, k_4P_, k_PIP5K_, k_5P_, k_PLC_, k_deg_ were estimated using Nelder-Mead optimization. Model fits are shown as cyan lines for normalized PI(4)P (i), PI(4,5)P_2_ (ii), and IP_3_ (iii). (**B**) Model validation using independent PI(4,5)P_2_ data not used in optimization. Using the same kinetic parameters from (**A**) with only PLC concentration adjusted to reflect receptor occupancy, the model predicted dose-dependent PI(4,5)P_2_ depletion at 1.0 μM (blue shading) and 10 μM Oxo-M (pink shading). Experimental PI(4,5)P_2_ measured via IKNCQ2/3 tail current (orange symbols in B-ii, Kruse et al., 2016) matched model predictions: 60% at 1.0 μM and 80-90% at 10 μM, validating the optimized parameters. Model predictions for PI(4)P (i), PI(4,5)P_2_ (ii), and IP_3_ (iii) are shown as black lines, with panel (iii) showing free IP_3_ (solid) and biosensor LIBRAvIII-bound IP_3_ (dashed).

**Table 1.** SCG neuron parameters were obtained by Nelder-Mead optimization (10 μM Oxo-M, 20s stimulation). Fingerprint and NN+SS estimates are shown for tsA201, neuroblastoma, and hippocampal cells. All parameters are first-order rate constants

|  |  | SCG Neurons | tsA201 Cells |  | Neuroblastoma Cells |  | Hippocampal Neurons |  |
| --- | --- | --- | --- | --- | --- | --- | --- | --- |
| Parameter | Units | Baseline | Fingerprint | NN + SS | Fingerprint | NN + SS | Fingerprint | NN + SS |
| k_PI4K | s<sup>-1</sup> | 0.000778 | 0.000474 | 0.000886 | 0.000158 | 0.000437 | 0.001106 | 0.000977 |
| k_4P | s<sup>-1</sup> | 0.040192 | 0.016590 | 0.031022 | 0.003542 | 0.009790 | 0.005950 | 0.005254 |
| k_PIP5K | s<sup>-1</sup> | 0.016261 | 0.009744 | 0.014247 | 0.022736 | 0.032593 | 0.001624 | 0.001467 |
| **k_5P** | $s^{-1}$ | **0.021756** | 0.013032 | 0.011398 | 0.030408 | 0.043685 | 0.021720 | 0.015895 |
| **k_PLC** | $s^{-1}$ | **0.241987** | 0.242000 | 0.257207 | 0.242000 | 0.254839 | 0.242000 | 0.170674 |
| **k_deg** | $s^{-1}$ | **0.066021** | 0.070000 | 0.061452 | 0.070000 | 0.059343 | 0.070000 | 0.084067 |
| *Initial conditions [PI(4)P□, PI(4,5)P□□, IP□, LIBRAVIII, IP□-LIBRAVIII]: SCG [4540, 3232, 0, 6, 0] \| tsA [4000, 5000, 0, 6, 0] \| Neuroblastoma [768, 573, 0, 6, 0] \| Hippocampal [26000, 2400, 0, 6, 0]* |  |  |  |  |  |  |  |  |

To validate the model, we tested dose-dependent PI(4,5)P_2_ responses using a different dataset not used in optimization (**Figure 2B**). Agonist concentrations were simulated by adjusting PLC activity levels (PLC_1_ = 0.15 for 1.0 μM and PLC_1_ = 0.7 for 10 μM Oxo-M) while holding all kinetic parameters constant. The model accurately reproduced experimentally observed PI(4,5)P_2_ depletion measured via IK(M) (KCNQ2/3) tail currents (Kruse et al., 2016) (**Figure 2B-ii**), with approximately 60% depletion at 1.0 μM Oxo-M and 80-90% depletion at 10 μM Oxo-M. The validation supports that six parameters are sufficient to capture the signaling in SCG neurons under different stimulation protocols. The model also captured dose-dependent IP_3_ dynamics (**Figure 2B-iii**), explicitly accounting for IP_3_ biosensor (LIBRAvIII) binding by distinguishing between free and biosensor-bound IP_3_ pools, reproducing both the magnitude and kinetics of IP_3_ accumulation and clearance across different agonist concentrations.

We next applied local and global PRCC sensitivity analysis to the optimized superior cervical ganglion (SCG) neuron model to map kinetic parameters to distinct dynamic signaling features. The ±20% parameter variation range was selected based on reported coefficients of variation in phosphoinositide pool sizes and kinase activity levels across individual cells of the same type, which typically fall within 15-25% (Kruse et al., 2016). This range reflects within-cell-type biological variability rather than cross-cell-type differences, and its appropriateness is supported by the observation that SCG neuron model variants within ±20% captured the experimental variability in that reference dataset. Sensitivity metrics were extracted for three experimentally observable features: minimum PI(4)P, PI(4,5)P_2_ recovery, and peak IP_3_* amplitude.

Local sensitivity analysis used a one-factor-at-a-time (OFAT) approach to quantify the isolated effect of each parameter, varying each parameter ±20% independently, while holding all other parameters constant. Minimum PI(4)P was affected by k_PI4K_ and k_4P_ (**Figure 3A-i**), with similar effects from k_PIP5K_ and k_5P_ (**Figure 3B-i**) but minimal effects due to k_PLC_ and k_deg_ (**Figure 3C-i**). PI(4,5)P_2_ recovery exhibited distributed sensitivity across k_PI4K_ and k_4P_ (**Figure 3A-ii**), k_PIP5K_ and k_5P_ (**Figure 3B-ii),** and to a lesser extent k_PLC_ and k_deg_ (**Figure 3C-ii**), indicating that recovery dynamics require coordinated variation across multiple enzymatic steps. IP_3_* peak amplitude and kinetic variations were masked by trace-wise normalization in panels **3A-C**(iii), but absolute sensitivity is quantified in panel D(iii). IP_3_* dynamics were primarily influenced by k_deg_ (**Figure 3C-iii**), which controlled IP_3_* decay kinetics. Quantitative OFAT sensitivity metrics for minimum PI(4)P, PI(4,5)P_2_ recovery, and peak IP_3_* amplitude are shown in **Figure 3D**, where paired bars reveal asymmetric parameter effects: increasing versus decreasing individual parameters produced different magnitudes of change, reflecting nonlinear pathway dynamics around the baseline operating point.

**Figure 3.**
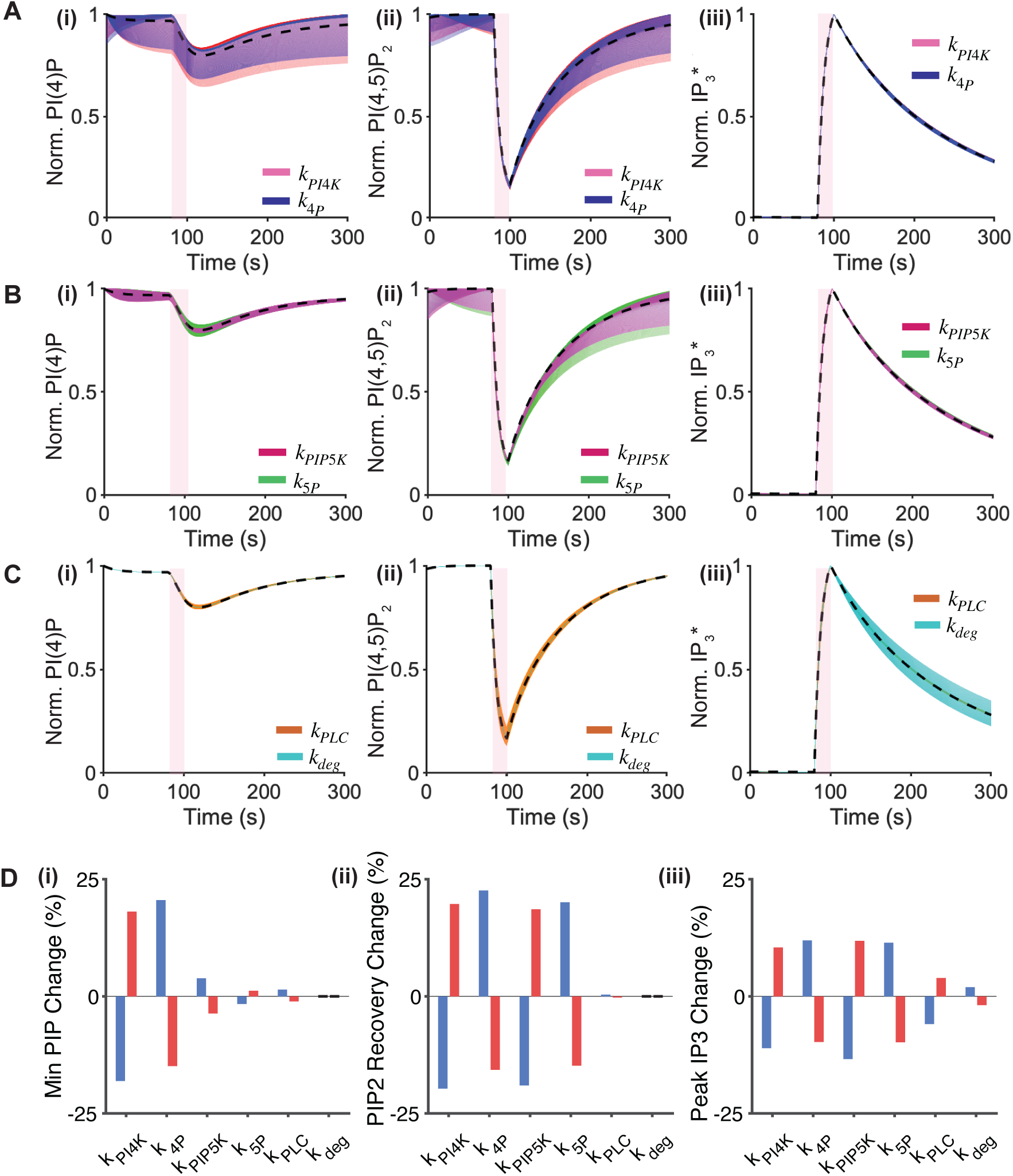
Local sensitivity analysis using the one-factor-at-a-time (OFAT) approach. (**A–C**) Local sensitivity analysis of the SCG baseline model from Figure 2A. Each parameter was varied independently by ±20% while holding all others constant, following 10 μM Oxo-M stimulation for 20s (pink shading). Parameters grouped by pathway step: k_PI4K_ and k_4P_ (**A**), k_PIP5K_ and k_5P_ (**B**), k_PLC_ and k_deg_ (**C**). Shaded regions show the range of responses for normalized PI(4)P (i), PI(4,5)P_2_ (ii), and IP_3_*(iii). IP_3_* represents biosensor-bound IP_3_ (IP_3_-LIBRAvIII), the experimentally measured quantity. Dashed black lines show the baseline. (**D**) Quantitative sensitivity metrics from OFAT analysis. For each parameter, two bars show the percent change in the metric when that parameter is decreased by 20% (blue bars) or increased by 20% (red bars) relative to baseline. Bar height indicates sensitivity magnitude. Positive bars indicate the parameter change increases the metric; negative bars indicate the parameter change decreases the metric. Metrics shown: minimum PI(4)P (**i**), PI(4,5)P_2_ recovery at t=300s (final/initial ratio) (**ii**), and peak of IP_3_ (**iii**).

Global Partial Rank Correlation Coefficients (PRCC) sensitivity analysis was performed by varying all six kinetic parameters simultaneously by ±20% using Latin Hypercube Sampling to generate 5,000 parameter sets. The model remained stable under combined parameter perturbations and generated a bounded population of signaling responses **(Figure 4A-i, ii, iii)**. PRCC quantified how each parameter influences signaling metrics when all parameters vary together (**Figure 4B**). For minimum PI(4)P **(Figure 4B-i)**, k_PI4K_ and k_PIP5K_ showed positive correlations, while k_PLC_ and k_4P_ showed negative correlations, confirming that PI(4)P levels depend on the balance between synthesis and degradation even under combined parameter variation. For PI(4,5)P_2_ recovery **(Figure 4B-ii)**, k_PI4K_ and k_PIP5K_ showed negative correlations, while all other parameters showed positive correlations, indicating that PI(4,5)P_2_ recovery dynamics require coordinated changes across multiple enzymatic steps. For IP_3_* production (**Figure 4B-iii**), k_PI4K_, k_PIP5K_, and k_PLC_ showed positive correlations as they drive IP_3_ generation through the synthesis pathway, while k_4P_, k_5P_, and k_deg_ showed negative correlations by opposing IP_3_ accumulation through substrate depletion or degradation. Together, local and global PRCC sensitivity analyses define a clear mapping between kinetic parameters and observable signaling features, with PRCC analysis confirming that parameter effects identified through OFAT persist under combined variation, providing a framework for interpreting cell-to-cell variability.

**Figure 4.**
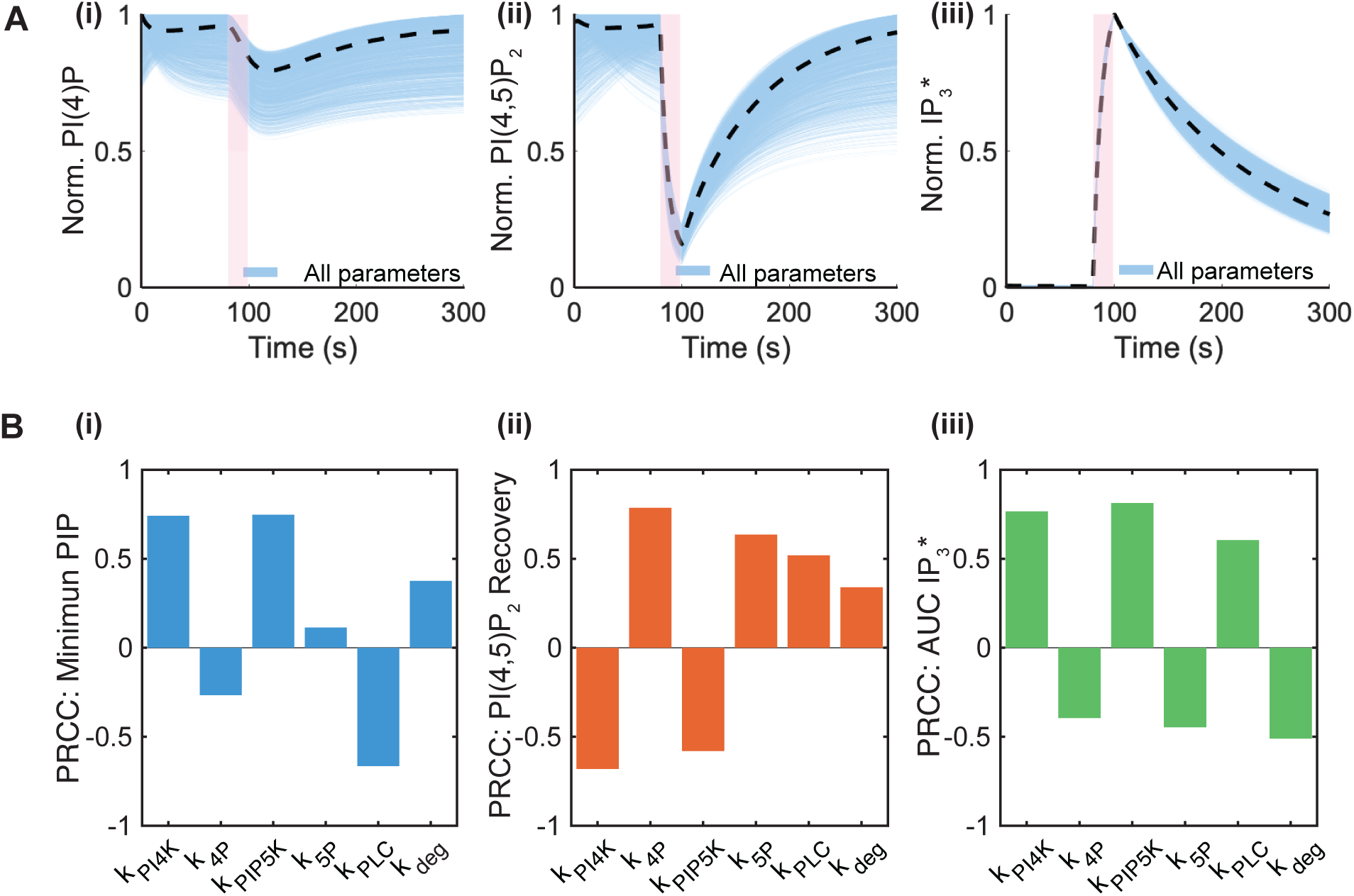
Global sensitivity analysis with all parameters varied simultaneously. (**A**) Model population generated by varying all six kinetic parameters simultaneously by ±20% using Latin Hypercube Sampling (N=5000 parameter sets). Shaded regions show the population envelope for normalized PI(4)P (i), PI(4,5)P_2_ (ii), and IP_3_*(iii) following 10 μM Oxo-M stimulation for 20s (pink shading). IP_3_* represents biosensor-bound IP_3_ (IP3-LIBRAvIII), the experimentally measured quantity. Dashed black lines show the baseline model. (**B**) Partial Rank Correlation Coefficients (PRCC) quantifying the strength and the influence of each parameter on signaling metrics when all parameters vary together. PRCC values range from -1 to +1, where magnitude indicates sensitivity strength and sign indicates direction of influence. Metrics shown: minimum PI(4)P (i), PI(4,5)P_2_ recovery (ii), and area under the curve (AUC) IP_3_ (iii).

We next tested whether sensitivity-constrained parameter variation could capture biological heterogeneity across cell types. Rather than refitting the model to each experimental dataset, we generated a population of model variants by varying each kinetic parameter by ±20% relative to the SCG baseline model. The ±20% range was selected to represent within-cell-type biological variability, providing a reference for identifying systematic cross-cell-type differences. Each model variant was evaluated using the stimulation protocol and experimental conditions specific to each cell type.

In SCG neurons, the model population reproduced the observed range of PI(4)P, PI(4,5)P_2_, and IP_3_* dynamics under 10 μM Oxo-M stimulation for 20s, establishing a reference distribution for identifying cell-type-specific parameter variation (**Figure 5A**). In tsA201 cells exposed to the same ligand and stimulation duration, experimental trajectories showed more PI(4)P and PI(4,5)P_2_ depletion than the SCG-derived model population (**Figure 5B**). These differences derive from parameters controlling lipid synthesis and turnover, including k_PI4K_, k_4P_, k_PIP5K_, and k_5P_. Additionally, the post-exposure PI(4,5)P_2_ recovery kinetics in tsA201 cells were slower than predicted by the SCG-derived population, likely reflecting differences in PI4K and PIP5K resynthesis rates between the two cell types (Dickson et al., 2013, Falkenburger et al., 2013). In human neuroblastoma cells stimulated with 1 mM carbachol for 60s, experimental responses showed deeper PI(4,5)P_2_ depletion and faster PI(4,5)P_2_ recovery than the SCG population (**Figure 5C**). These dynamics extended beyond the population distribution, indicating parameter variations exceeding the ±20% range. The range of variation was expanded empirically for each cell type based on the degree of deviation from the SCG-derived population. In hippocampal neurons stimulated with 10 μM L-glutamate for 50s, PI(4)P remained stable with no depletion due to larger basal PI(4)P pools, while PI(4,5)P_2_ and IP_3_* dynamics were captured by the SCG population (**Figure 5D**). The absence of PI(4)P depletion in hippocampal neurons is consistent with their larger basal PI(4)P pool, which confers greater buffering capacity (de la Cruz et al., 2022). Across all cell types, sensitivity-constrained population modeling revealed structured and feature-specific differences, highlighting specific enzymatic steps that vary across cellular contexts.

**Figure 5.**
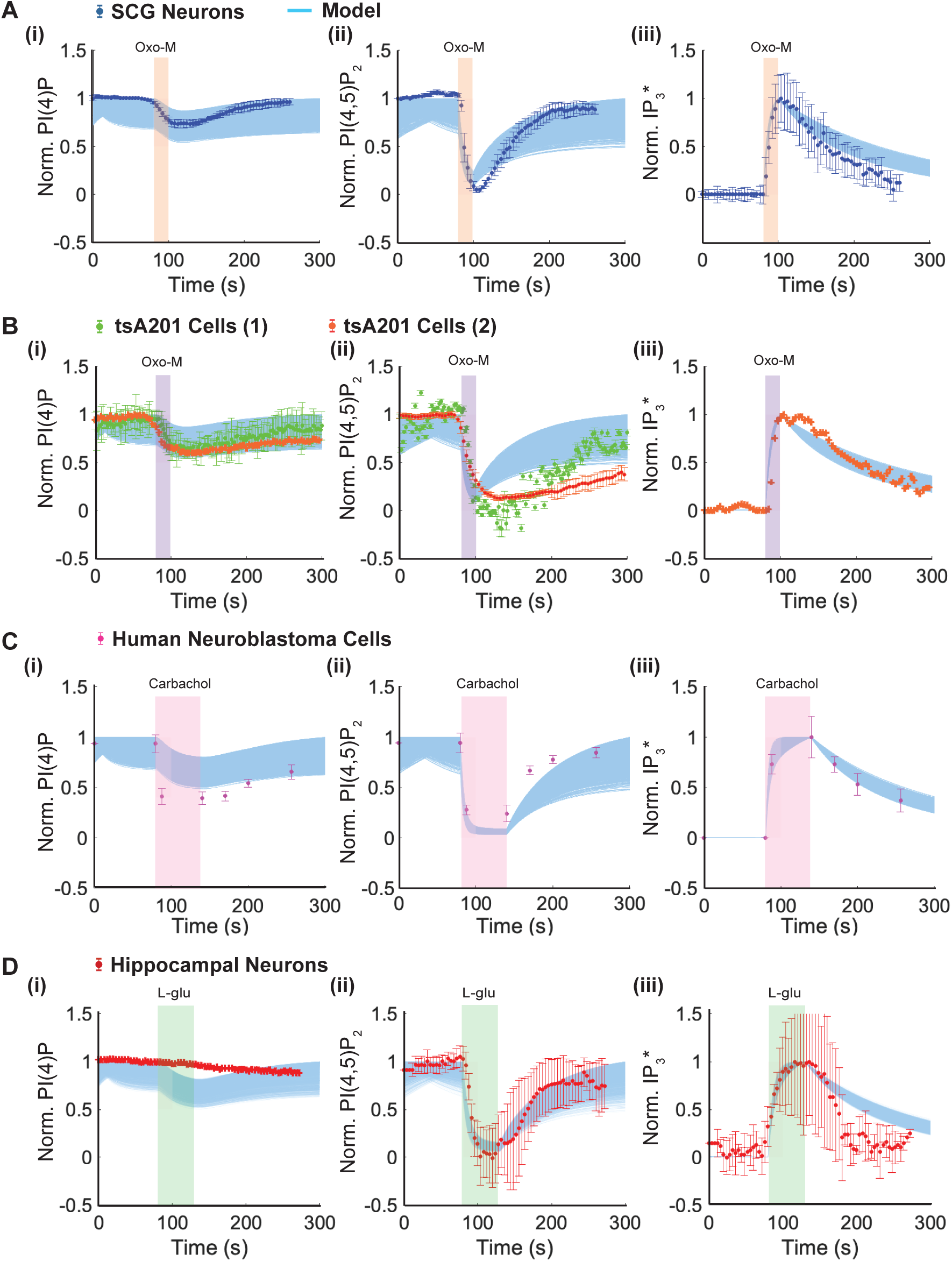
A population of models generated by varying model parameters captures phosphoinositide lipid dynamics in multiple cell types. A population of model variants was generated by perturbing kinetic parameters by ±20% in the SCG baseline neuron model shown in Figure 2A. (**A**) PI(4)P (i), PI(4,5)P_2_ (ii), and IP_3_*(iii) dynamics in SCG neurons following 10 μM Oxo-M stimulation for 20s (orange bar). Experimental data (symbols) shown with the model population (shaded region). (**B**) PI(4)P (i), PI(4,5)P_2_ (ii), IP_3_* (iii) dynamics in tsA201 cells following 10 μM Oxo-M stimulation for 20s (purple bar). Experimental data from two independent datasets (symbols) compared to the SCG-derived model population (shaded region). (**C**) PI(4)P (i), PI(4,5)P_2_ (ii), and IP_3_(iii) dynamics in human neuroblastoma cells following 1 mM carbachol stimulation for 60s (pink bar). Experimental data (symbols) compared to the SCG-derived model population (shaded region) using the corresponding stimulation protocol. (**D**) PI(4)P (i), PI(4,5)P_2_ (ii), and IP_3_* (iii) dynamics in hippocampal neurons following 10 μM L-glutamate stimulation for 50s (green bar). Experimental data (symbols) compared to the SCG-derived model population (shaded region) using the corresponding stimulation protocol. IP_3_* represents biosensor-bound IP_3_ (IP_3_-LIBRAvIII), the experimentally measured quantity (panels **A**(iii), **B**(iii), **D**(iii)). Panel **C**(iii) shows IP_3_ measured by a different method.

We next asked whether sensitivity structure could guide explicit parameter inference from experimental data. Guided by local sensitivity analysis from **Figure 3** and steady-state basal lipid pool compositions, we adjusted parameters for tsA201, neuroblastoma, and hippocampal models to generate cell-type-specific parameter fingerprints (**Figure 6A**). We then trained a hybrid convolutional neural network-long short-term memory (CNN-LSTM) neural network to map phosphoinositide time series to kinetic parameters. The broader ±40-60% parameter range used for neural network training was chosen to span cross-cell-type differences rather than within-cell-type variability, ensuring robust parameter recovery across the full range of dynamics observed experimentally. **Figure 6B** shows the neural network-predicted parameter changes for each cell type, which were consistent with the manually derived fingerprints. When applied to experimental measurements, both approaches inferred parameter sets that reproduced observed PI(4)P, PI(4,5)P_2_, and IP_3_* dynamics across cell types (**Figure 6C–E**). Inferred parameters were consistent across methods, particularly for k_PI4K_, k_4P_, and k_PIP5K_, which accounted for much of the observed cell-type variability. Together, sensitivity-constrained inference enabled bidirectional mapping between phosphoinositide dynamics and kinetic parameters, supporting direct construction of cell-type-specific signaling models from experimental measurements (**Table 1**).

**Figure 6.**
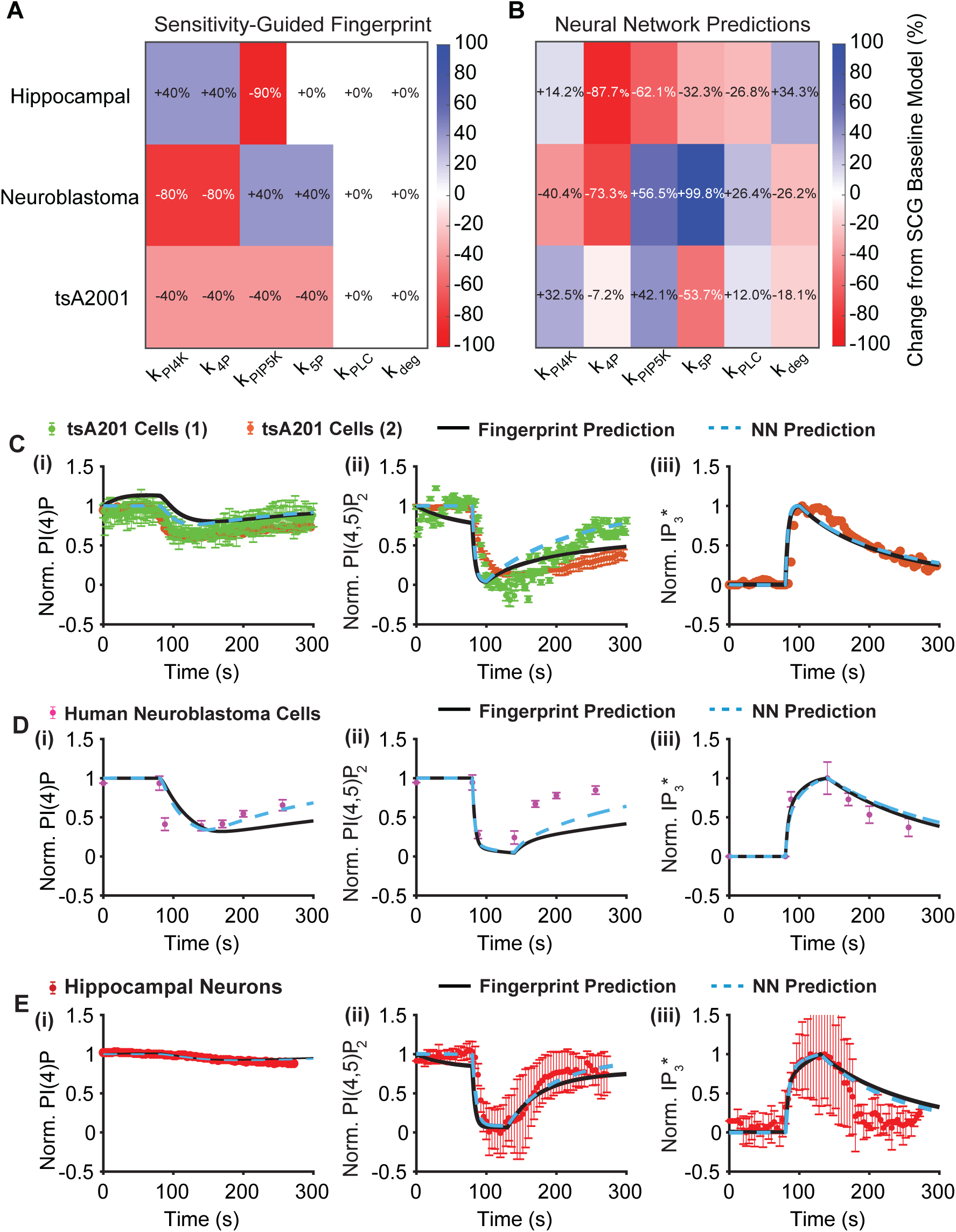
Parameter inference via sensitivity fingerprinting and neural network inverse modeling. (**A**) Sensitivity-informed parameter fingerprints for each cell type relative to the SCG baseline model from Figure 2A, based on sensitivity relationships (Figure 3) and observed deviations from SCG dynamics. (**B**) Neural network parameter predictions for each cell type relative to the SCG baseline model. The network was trained on synthetic phosphoinositide time series generated by varying SCG baseline parameters ±40-60%, with the known PI/PI(4)P ratio supplied as a fixed input channel. k_PIP5K_ was analytically adjusted post-inference to enforce pre-stimulus steady-state. (**C-E**) Model predictions using fingerprint-derived (black lines) versus neural network-inferred (cyan dashed lines) parameters compared to experimental data (symbols). (**C**) tsA201 cells, two independent datasets. (**D**) Neuroblastoma cells. (**E**) Hippocampal neurons. IP_3_* (panels **C**(iii) and **E**(iii)) represents biosensor-bound IP_3_ (IP_3_-LIBRAvIII).

Having established that cell-type-specific parameter variations could be systematically identified and quantified, we next asked whether this framework could predict signaling perturbations associated with loss-of-function mutations in key phosphoinositide enzymes. We simulated loss-of-function perturbations in PI4KA and PIP5K1C, genes linked to neurodevelopmental disorders with altered phosphoinositide homeostasis, by reducing k_PI4K_ and k_PIP5K_, respectively, and evaluated signaling responses under repeated receptor stimulation.

Simulations consisted of three 20-second stimulation pulses delivered at 100s, 600s, and 1100s. In superior cervical ganglion neurons, PI4KA loss-of-function was simulated by reducing k_PI4K_ by 50% and 100%. Under repeated stimulation, PI4K reduction caused progressive depletion of PI(4)P and PI(4,5)P_2_. Recovery failed between stimulation pulses as limited lipid reserves became exhausted. IP_3_ responses decreased with each pulse, indicating cumulative pathway failure (**Figure 7A**).

**FIGURE 7.**
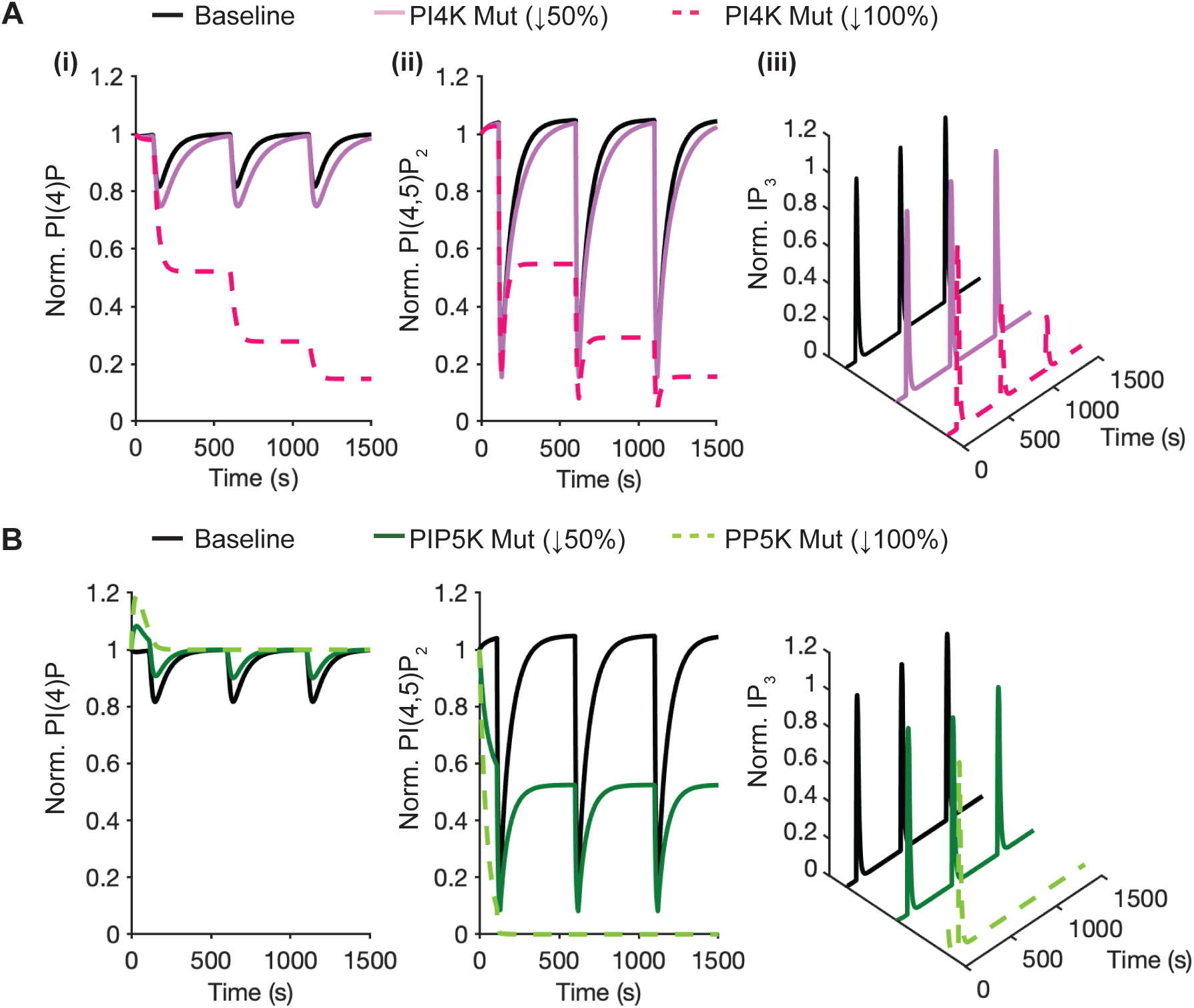
PI4K and PIP5K mutations disrupt lipid signaling in SCG neurons. Three 20s stimulation pulses (at 100s, 600s, and 1100s) were applied to the SCG baseline model from Figure 2A. Model predictions shown for PI(4)P (i), PI(4,5)P_2_ (ii), and IP_3_ (iii) dynamics. Black lines show the baseline. (**A**) PI4KA loss-of-function simulated by reducing k_PI4K_. Purple solid line: 50% reduction; pink dashed line: 100% reduction. Progressive PI(4)P and PI(4,5)P_2_ depletion with 100% reduction, showing failed recovery between pulses. (**B**) PIP5K1C loss-of-function simulated by reducing k_PIP5K_. Green solid line: 50% reduction; green dashed line: 100% reduction. Progressive reduction of PI(4,5)P_2_ synthesis, preventing PI(4,5)P_2_ recovery at 100% inhibition.

PIP5K1C loss-of-function was simulated by reducing kPIP5K by 50% and 100%, representing partial enzymatic impairment and complete functional loss, respectively. PIP5K reduction disrupted PI(4,5)P2 synthesis, leading to accumulation of PI(4)P before stimulation and depletion of PI(4,5)P_2_. IP_3_ production continued only while existing PI(4,5)P_2_ reserves remained available. Once depleted, IP_3_ responses collapsed during subsequent stimulation pulses. In SCG neurons, which maintain small basal PI(4)P pools, both PI4K and PIP5K perturbations produced progressive signaling failure under repeated stimulation (**Figure 7B**).

We next examined whether the same mutations produced different outcomes in cell types with larger basal PI(4)P pools. Using the same stimulation protocol with 20s pulses at 100s, 600s, and 1100s, simulations were performed in hippocampal neurons, which maintain substantially larger PI(4)P pools than SCG neurons (26,000 vs 4,540 molecules/μm², a 5.7-fold difference)(Kruse et al., 2016). In the hippocampal context, k_PI4K_ reduction produced a markedly different response. PI(4)P and PI(4,5)P_2_ recovered between stimulation pulses under both cases of 50% and 100% reduction. IP_3_ signaling was preserved across all three pulses, indicating that large basal PI(4)P reserves buffer the effects of reduced PI4K activity (**Figure 8A**).

**Figure 8.**
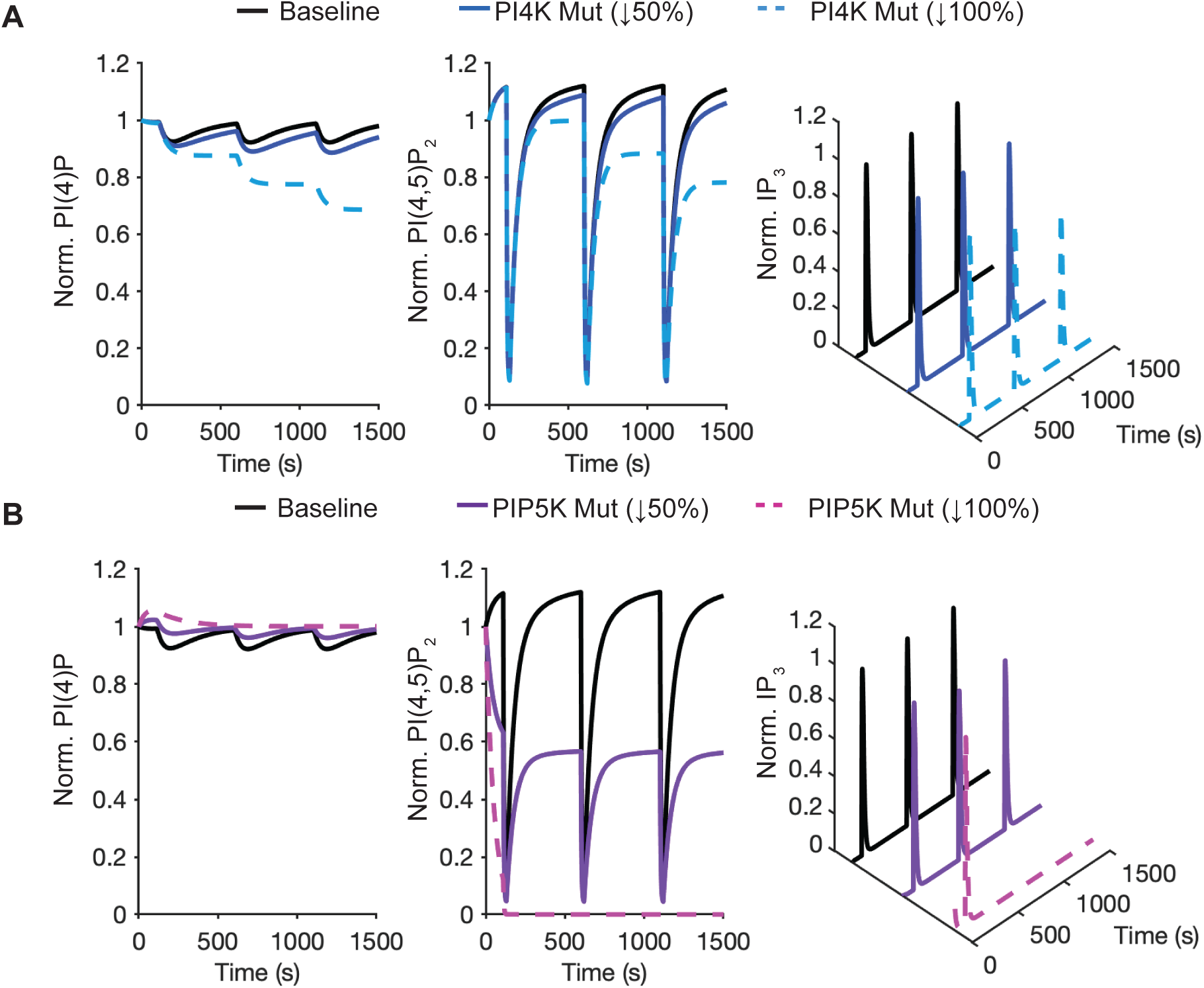
Hippocampal neurons with larger PI(4)P lipid pools are resistant to PI4K mutations. Three 20s stimulation pulses (at 100s, 600s, and 1100s) were applied to the hippocampal neuron model. Hippocampal neurons have much larger basal PI(4)P pools (15.4%) compared to SCG neurons (1.9% in Figure 7). Model predictions for PI(4)P (i), PI(4,5)P_2_ (ii), and IP_3_ (iii). Black: baseline. (**A**) PI4KA loss-of-function simulated by reducing k_PI4K_. Blue solid: 50% reduction; light blue dashed: 100% reduction. PI(4)P and PI(4,5)P_2_ recover between pulses despite 100% mutation. IP_3_ signaling was maintained across all three pulses. Larger PI(4)P pools protect against the same mutation that caused failure in SCG neurons (Figure 7A). (**B**) PIP5K1C loss-of-function simulated by reducing k_PIP5K_. Purple solid: 50% reduction; magenta dashed: 100% reduction. PI(4,5)P_2_ depletion is similar to that of SCG neurons. IP_3_ responses sustained longer due to larger baseline pools. Larger lipid pools provide partial protection, though 100% enzyme loss eventually prevents PI(4,5)P_2_ recovery.

PIP5K reduction in hippocampal neurons produced milder phenotypes compared to SCG neurons. Although loss of PIP5K activity ultimately prevented full PI(4,5)P_2_ restoration, larger baseline PI(4,5)P_2_ pools sustained IP_3_ responses for longer durations. PI(4,5)P_2_ depletion occurred more gradually, and IP_3_ signaling persisted across repeated stimulation before eventual decline. The simulations demonstrate that larger basal PI(4)P pools confer protection against genetic perturbations that cause pathway failure in cells with smaller lipid reserves (**Figure 8B**)

Together, the simulations demonstrate that the effects of lipid signaling mutations depend on the size of basal phosphoinositide pools. The same genetic perturbation causes severe signaling failure in cells with small lipid reserves but allows partial compensation in cells with larger reserves. By combining sensitivity analysis with kinetic modeling, the sensitivity-guided NAM links mutation effects to cellular biochemical context and enables the prediction of cell-specific vulnerability to signal disruption under genetic perturbation.

## Discussion

Here, we present a computational framework that integrates mechanistic modeling, sensitivity analysis, and parameter inference to predict cell-specific signaling behavior, which together comprise a novel NAM (new approach methodology). Using phosphoinositide signaling as a proof of concept, this work demonstrates how dynamic biochemical measurements can be translated into cell-specific predictive models, provide new disease insights, and serve as a potential tool for precision medicine.

We first validated a kinetic model of PI, PI(4)P, PI(4,5)P_2_, and IP_3_ signaling (**Figure 1**). The model includes only the enzymatic steps required to reproduce experimentally measured dynamics and conserves lipid mass across pathway components. Kinetic parameters were optimized and validated using independent experimental datasets from superior cervical ganglion neurons (**Figure 2A, B**). Agreement between model predictions and experimental measurements provided a stable and biologically grounded foundation for subsequent analyses.

Local and global PRCC sensitivity analysis revealed how kinetic parameters subtly control different aspects of signaling, rather than a dominant rate-limiting step (**Figures 3 and 4**). PI(4)P depletion, PI(4,5)P_2_ recovery, and IP_3_ production were governed by overlapping but distinct subsets of rate constants (**Figure 3A-E**). Building on this structure, we tested whether sensitivity analysis could guide inference of cell-specific parameters from experimental data. Bounded parameter variation (±20%) generated a population of models that captured phosphoinositide dynamics across multiple cell types without refitting the model (**Figure 5**).

We demonstrated two approaches to determine rate parameters in the model (**Figure 6**). One method used sensitivity analysis to identify the most important parameters that needed to be adjusted to match the data. The other method used a neural network to automatically estimate rates from the data. Both methods were able to closely match the measured changes over time in PI(4)P, PI(4,5)P_2_, and IP_3_ in three different cell types (tsA201 cells, neuroblastoma cells, and hippocampal neurons). Because the two approaches gave similar results, this suggests that time-based measurements of lipid signaling contain enough information to uncover real differences in the underlying biology, assuming that model structure is plausible. The convergence of the two methods also provides practical evidence of parameter identifiability: when independent approaches starting from different assumptions recover consistent parameter sets, it reduces concern that the data are compatible with a broad family of degenerate solutions. Formal identifiability analysis remains an important caveat, however. The six-parameter model is deliberately parsimonious, and some parameter combinations that control similar observables (for example, k_PI4K_ and k_4P_ both influence minimum PI(4)P) may be partially compensatory. We note that sensitivity analysis explicitly maps which parameters are distinguishable through specific observables, providing a practical guide to which parameters can be reliably inferred from a given dataset. For the neural network inference method, the experimentally measured PI/PI(4)P ratio for each cell type was supplied as a fixed input channel, and network predictions were refined by steady-state correction to ensure biologically consistent pre-stimulus conditions.

We next simulated disease-relevant perturbations in PI4KA and PIP5K1C (**Figures 7 and 8**) by implementing variants as a reduction in k_PI4K_ or k_PIP5K_, to predict how naturally occurring human genetic mutations may alter lipid synthesis by reducing enzyme catalytic activity or lipid abundance, or by impairing membrane recruitment or protein complex stability. PI4KA encodes PI4KIIIα, which generates the major plasma-membrane PI4P pool that supplies PI(4,5)P_2_ synthesis and is regulated through assembly and recruitment of a PI4KIIIα–TTC7–FAM126 complex(Nakatsu et al., 2012; Suresh et al., 2024). Importantly, experimental perturbation of PI4KA shows that it is required to maintain plasma-membrane PI(4,5)P_2_ during strong PLC-coupled receptor stimulation.

A key prediction from the model simulations indicates the role of basal lipid pool composition in determining genetic vulnerability. Under repeated stimulation, cells with small basal PI(4)P pools exhibited progressive depletion of PI(4)P and PI(4,5)P_2_ and collapse of IP_3_ signaling when PI4KA or PIP5K1C flux was reduced (**Figure 7**), whereas hippocampal neurons with larger basal PI(4)P pools retained partial recovery and sustained signaling even under severe perturbation (**Figure 8**). The fragile versus protected behavior is consistent with experimentally observed responses to kinase perturbation. Hippocampal neurons, which maintain large PI(4)P pools (26,000 vs 4,540 molecules per square micrometer in SCG neurons), recover PI(4,5)P_2_ normally even when PI4K is pharmacologically inhibited, whereas cells with small PI(4)P pools cannot recover without active PI4K (de la Cruz et al., 2022). Our simulations reproduce the behavior: reducing k_PI4K_ by 50 to 100% prevents PI(4,5)P_2_ recovery in SCG neurons but has minimal effect on hippocampal neurons (**Figures 7A-ii vs 8A-ii**).

The results provide a mechanistic explanation for tissue-specific disease severity observed in disorders linked to PI4KA and PIP5K1C mutations (Burke et al., 2023; Narkis et al., 2007; Salter et al., 2021; Verdura et al., 2021) . While the simulations presented here focus on loss-of-function mutations, PIP5K1C gain-of-function variants have also been reported and are associated with excessive IP_3_ production and pathological calcium signaling (Morleo et al., 2023). Extension of the framework to simulate gain-of-function conditions suggests uncontrolled IP_3_ generation as a potential mechanism for disrupted signaling and cellular dysfunction.

One future opportunity is to address the limitation that the model presented herein does not explicitly account for the energetic regulation of kinase activity. Because plasma-membrane PI(4)P and PI(4,5)P_2_ can be depleted rapidly under hypoxia and ATP inhibition, the energetic state represents an additional axis of biochemical regulation captured by k_PI4K_ and k_PIP5K_ (Lu et al., 2022). In excitable tissues, ATP can fluctuate on rapid timescales (including beat-resolved ATP dynamics in cardiac myocytes), raising the possibility that ATP microdomains modulate phosphoinositide resynthesis capacity during repeated demand (Rhana et al., 2024). Future extensions incorporating energetic constraints would complement established lipid-homeostatic feedback, such as PIP4K-dependent restraint of PI(4,5)P_2_ synthesis by PIP5Ks (Wang et al., 2019; Wills et al., 2023).

Together, the work demonstrates a New Approach Methodology (NAM) in which mechanistic modeling, sensitivity analysis, and constrained inference translate dynamic lipid measurements into interpretable, cell-specific kinetic parameters for modeling and simulation. In a precision-medicine context, the framework naturally supports a workflow in which a patient genotype is paired with time-resolved PI, PI(4)P, PI(4,5)P_2_, and IP_3_ measurements in patient-derived cells to infer effective changes resulting from genetic perturbation. The arrival of increasingly sophisticated biosensors makes this plausible in the not-too-distant future. More broadly, the increasing therapeutic interest in phosphoinositide kinases underscores the value of quantitative, context-aware models for connecting variant mechanisms to predicted signaling phenotypes.

## Methods

### Phosphoinositide Signaling Model Structure

A system of differential equations was developed to represent the dynamic interconversion of phosphatidylinositol (PI), phosphatidylinositol 4-phosphate (PI(4)P), and phosphatidylinositol 4,5-bisphosphate (PI(4,5)P_2_), as well as the hydrolytic cleavage of PI(4,5)P2 by phospholipase C (PLC), yielding inositol 1,4,5-trisphosphate (IP_3_). The model structure was adapted and simplified from a previously published formulation describing the signaling dynamics in tsA-201 cells and superior cervical ganglion (SCG) neurons(Falkenburger et al., 2013; Kruse et al., 2016) . The model tracks five dynamic state variables: PI(4)P and PI(4,5)P_2_ on the plasma membrane (molecules/μm²), free cytoplasmic IP_3_ (μM), and the genetically encoded IP_3_ biosensor LIBRAvIII in both free and IP_3_-bound states (μM). PI was treated as a constant pool because PI accounts for more than 90 percent of the total phosphoinositide pool in both superior cervical ganglion neurons and tsA 201 cells (Kruse et al., 2016).

The governing equations for membrane species (molecules/μm²) are:

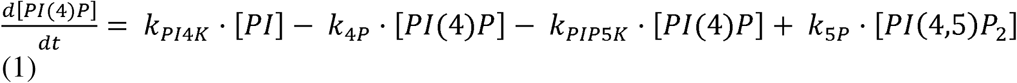

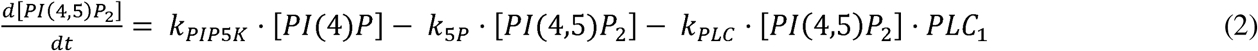

For cytoplasmic species (μM), the equations describing IP_3_ dynamics and biosensor binding are:

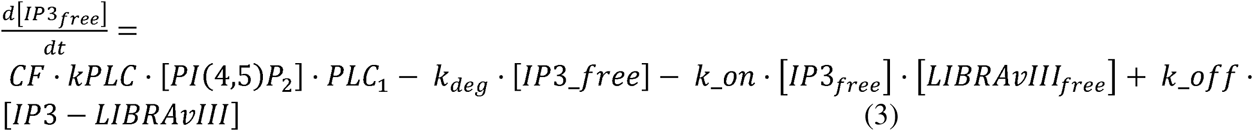

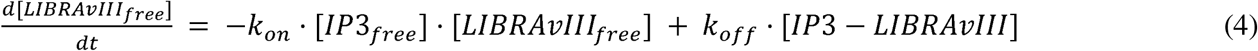

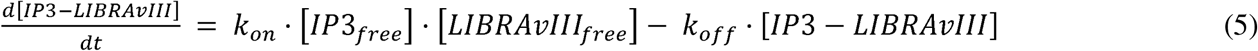

where the biosensor kinetics parameters *k*_on_ = 1.0 s^−1^ and *k*_off_ = 0.5 s^−1^, were obtained experimentally, yielding an equilibrium dissociation constant *K*_D_ = 0.5 μM (Kruse et al., 2016). The conversion factor CF links membrane PI(4,5)P_2_ hydrolysis to cytoplasmic IP_3_ production: CF = (*S*_A_ / *V*_cyt_) × (1/602) = (4100 μm² / 6644 μm³) × (1/602) = 1.03 × 10^−3^ μm^−1^, where *S*_A_ is the plasma membrane surface area, *V*_cyt_ is the cytoplasmic volume, and 1/602 converts molecules to μM·μm³ (Kruse et al., 2016).

Receptor activation by muscarinic agonist (10 μM oxotremorine-M) was modeled as a step function: PLC_1_ = 0.7 (dimensionless activity coefficient) during stimulation and PLC_1_ = 0 otherwise. For validation data (Figure 2B), PLC_1_ = 0.15 and PLC_1_ = 0.7 were used to simulate application of 1.0 μM and 10.0 μM oxotremorine-M, respectively.

Initial conditions at *t* = 0 were set to experimentally measured basal levels: [PI(4)P]_0_ = 4540 molecules/μm², [PI(4,5)P_2_]_0_ = 3232 molecules/μm², [IP_3free_]_0_ = 0.02 μM, [LIBRAvIII_free_]_0_ = 6.0 μM, and [IP3-LIBRAvIII]_0_ = 0.0 μM. The constant PI pool was fixed at [PI] = 226,975 molecules/μm² (Kruse et al., 2016).

The system of five coupled ODEs was solved numerically over *t* ∈ [0, 300] s using the MATLAB ordinary differential equation (ode15s) solver with relative tolerance 10^−6^ and absolute tolerance 10^−10^. Model outputs were compared to experimental measurements of the PI(4)P, PI(4,5)P_2_, and IP_3_-LIBRAvIII biosensor signal.

### Nelder-Mead Parameter Optimization of the Phosphoinositide Signaling Model

The Nelder-Mead method was used to estimate the kinetic parameters governing the time-dependent behavior of the phosphoinositide intermediates PI(4)P, PI(4,5)P_2_, and IP_3_. The system comprises six parameters (k_PI4K_, k_4P_, k_PIP5K_, k_5P_, k_PLC_, k_deg_). Nelder-Mead is a derivative-free optimization method well-suited for biological parameter estimation problems where the cost function landscape contains multiple local minima (Hernandez-Hernandez et al., 2024; Kernik et al., 2019).

The cost function for optimization was defined as the total sum of squared errors (SSE) across all time points:

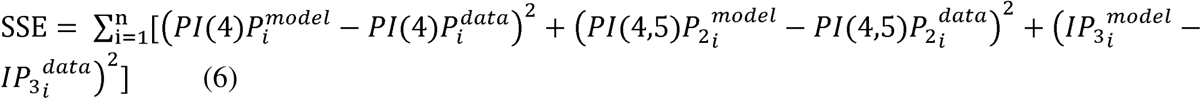

A bounded transformation confined all parameters to the biologically plausible range of 10 to 1000. Convergence was declared when both the maximum relative change in all parameters fell below 10 and the relative change in SSE fell below 10 between consecutive iterations, with a maximum of 10,000 function evaluations per trial.

To mitigate the risk of convergence to local minima, a multi-start global optimization strategy was employed. Each trial was initialized with a randomly generated parameter vector sampled from log-uniform distributions spanning the allowed parameter range. Upon identifying a local minimum, the parameters were perturbed by independently adjusting each value within ±20%, and the resulting vector was used to initiate a new optimization run. This process was repeated for 1,000 independent trials. The parameter set yielding the lowest total sum of squared errors (SSE) across all iterations was selected as the optimal solution.

Model accuracy was assessed through residual diagnostics. Metrics included the sum of squared errors, the root mean square error, as well as error decomposition by molecular species.

### Local Sensitivity Analysis

We performed one-factor-at-a-time (OFAT) local sensitivity analysis to quantify how individual parameter perturbations affect model outputs. For each of the six kinetic parameters (k_PI4K_, k_4P_, k_PIP5K_, k_5P_, k_PLC_, k_deg_), we generated 5,000 model simulations spanning parameter values from 0.8× to 1.2× baseline. The ±20% variation range reflects the inter-replicate variability observed in the SCG neuron optimization dataset (Kruse et al., 2016) and is consistent with reported coefficients of variation in phosphoinositide pool sizes and kinase expression levels across individual cells of the same type, which typically fall within 15–25% (de la Cruz et al.,2022). All other parameters were held constant at baseline values during each parameter variation.

Three metrics were calculated from each simulation: (1) minimum PI(4)P, (2) PI(4,5)P_2_ recovery at t=300s (final/initial ratio), and (3) peak IP3 amplitude. Stimulation was 10 μM Oxo-M applied from 80s to 100s.

### Global Sensitivity Analysis

To assess parameter interactions and system robustness beyond one-factor-at-a-time (OFAT) analysis, we performed global sensitivity analysis by simultaneously varying all six kinetic parameters (k_PI4K_, k_4P_, k_PIP5K_, k_5P_, k_PLC_, k_deg_), each sampled uniformly in log-space across the ±20% range used for local analysis. We generated 5,000 parameter sets using Latin Hypercube Sampling (LHS), a space-filling quasi-random sampling method that ensures efficient coverage of multidimensional parameter space by dividing each dimension into N equal-probability intervals, sampling one value per interval, then permuting samples across dimensions to minimize correlation.

To quantify parameter importance in the global analysis, we calculated Partial Rank Correlation Coefficients (PRCCs) between each parameter and output metrics to measure monotonic association strength after removing linear effects of all other parameters. PRCC values range from −1 to +1, where values near ±1 indicate strong monotonic relationships and values near 0 indicate weak or complex non-monotonic effects.

### Neural Network Parameter Estimation

To enable rapid parameter estimation from experimental time-series data, we trained a hybrid convolutional neural network-long short-term memory (CNN-LSTM) neural network to predict kinetic parameters from normalized PI(4)P, PI(4,5)P2, and IP_3_ trajectories. Convolutional layers extract local temporal features while LSTM layers capture long-range sequential dependencies across the full-time course. The network comprised two 1D convolutional layers (64 and 128 filters, kernel size 5) with batch normalization and max pooling for temporal feature extraction, followed by a 2-layer LSTM (128 hidden units) to capture sequential dynamics, and three fully connected layers (128, 64, 5 units) for parameter prediction. The experimentally measured PI/PI(4)P ratio for each cell type, derived from basal lipid pool compositions, was supplied as a fixed fifth input channel rather than treated as a predicted output. The network predicted five kinetic parameters (k_PI4K_, k_PIP5K_, k_5P_, k_PLC_, k_deg_), with k_4P_ derived analytically as k_4P_ = (PI/PI(4)P ratio) × kPI4K.

Training data consisted of 50,000 simulated trajectories generated using Latin Hypercube Sampling across parameter ranges spanning ±40–60% of baseline values for the five kinetic parameters, with the PI/PI(4)P ratio sampled uniformly from 3 to 50. This range captured parameter variations observed in cell types beyond SCG neurons, particularly hippocampal neurons and neuroblastoma cells. Variable stimulus durations (20, 50, or 60s) matching experimental conditions were encoded as an additional input channel. Each simulation trajectory was interpolated to 100 time points over 300s, with each species (PI(4)P, PI(4,5)P2, IP_3_) normalized independently to its maximum value within that trajectory and paired with its corresponding parameter set as ground truth.

Training used log10-transformed parameters with mean squared error loss, Adam optimizer (learning rate 0.001, weight decay 10^−5^), batch size 64, and learning rate scheduling using a 90/10 train-validation split. Training proceeded up to 50 epochs with early stopping (patience = 10 epochs) to prevent overfitting. The model was implemented in PyTorch and trained on GPU when available.

For experimental data prediction, normalized PI(4)P, PI(4,5)P_2_, and IP_3_ time courses were input to the trained network along with stimulus duration. Network outputs in log_10_ space were converted back to linear scale via *k_i_* = 10*^yi^*, where *y_i_*is the network output for parameter *i*. Predicted parameters were compared to baseline SCG values as percentage changes. k_PIP5K_ was analytically adjusted to enforce pre-stimulus steady state by setting k_PIP5K_ = k5P × (PI(4,5)P_2,0_ / PI(4)P_0_), where PI(4,5)P_2,0_ and PI(4)P_0_ are the experimentally measured basal concentrations for each cell type. To assess reproducibility, the network was trained three independent times using different random seeds (42, 123, 456), and the mean predicted parameters across runs were used as the final cell-type-specific estimates.

### Simulation protocols

Code for simulations and analysis was written in MATLAB 2018b. The MATLAB optimization routine was run on an Apple Mac Pro machine with two 2.7 GHz 12-Core Intel Xeon processors and an HP ProLiant DL585 G7 server with a 2.7 GHz 28-core AMD Opteron processor. Numerical results were visualized using MATLAB R2018a by The MathWorks, Inc. All codes are available on GitHub (https://github.com/ClancyLabUCD/from-lipid-dynamics-to-precision-predictions)

## Declaration of generative AI and AI-assisted technologies in the manuscript preparation process

During the preparation of this work, the authors used ChatGTP and Claude for language editing purposes. After using these tools, the authors reviewed and edited the content as needed and take full responsibility for the content of the published article.

## Additional Information

### Competing Interests

The authors declare no competing interests.

### Funding

This work was supported by the National Heart, Lung, and Blood Institute (R01HL174001 and R01HL168874 to C.E.C. and L.F.S.), the National Institute of General Medical Sciences (R35GM142690 to O.V.), the UC Davis Chancellor’s Postdoctoral Fellowship (G.H.-H.), and the UC Davis School of Medicine Center for Precision Medicine and Data Sciences.

### Author Contributions (CRediT)

Conceptualization: Gonzalo Hernandez-Hernandez, Pei-Chi Yang, Colleen E. Clancy

Methodology: Gonzalo Hernandez-Hernandez, Pei-Chi Yang, Timothy J. Lewis, Colleen E. Clancy

Software: Gonzalo Hernandez-Hernandez

Validation: Gonzalo Hernandez-Hernandez, Pei-Chi Yang, Oscar Vivas

Formal Analysis: Gonzalo Hernandez-Hernandez, Mindy Tieu, Pei-Chi Yang, Oscar Vivas, Colleen E. Clancy

Investigation: Gonzalo Hernandez-Hernandez

Resources: Oscar Vivas, L. Fernando Santana, Colleen E. Clancy

Data Curation: Gonzalo Hernandez-Hernandez

Writing – Original Draft: Gonzalo Hernandez-Hernandez, Colleen E. Clancy

Writing – Review & Editing: Gonzalo Hernandez-Hernandez, Mindy Tieu, Pei-Chi Yang, Oscar Vivas, Timothy J. Lewis, L. Fernando Santana, Colleen E. Clancy

Visualization: Gonzalo Hernandez-Hernandez

Supervision: Colleen E. Clancy

Project Administration: Colleen E. Clancy

Funding Acquisition: Colleen E. Clancy, L. Fernando Santana, Oscar Vivas

## Acknowledgements

The authors thank Hannah Zukowski for valuable discussions and feedback during the development of this work.

## Data Availability Statement

All data supporting the findings of this study are contained within the article and its Supplementary Information. The MATLAB source code used for model development, parameter optimization, sensitivity analysis, neural network training, and simulation is publicly available at https://github.com/ClancyLabUCD/from-lipid-dynamics-to-precision-predictions. Experimental datasets used for model development were obtained from previously published studies, which are cited throughout the manuscript and listed in the References. Additional materials required to reproduce the analyses are available from the corresponding author upon reasonable request.

## Notes

https://github.com/ClancyLabUCD/from-lipid-dynamics-to-precision-predictions

### Competing Interest Statement

The authors have declared no competing interest.

### Summary of Updates

This version contains revisions throughout the manuscript, including improvements to the text, figures, analyses, and overall organization to enhance clarity, rigor, and presentation.

## References

Balla, T. (2013). Phosphoinositides: Tiny lipids with giant impact on cell regulation. Physiological Reviews, 93(3), 1019–1137. 10.1152/physrev.00028.2012

Berridge, M. J. (1993). Inositol trisphosphate and calcium signalling. Nature, 361(6410), 315–325. 10.1038/361315a0

Berridge, M. J. (2009). Inositol trisphosphate and calcium signalling mechanisms. Biochimica Et Biophysica Acta, 1793(6), 933–940. 10.1016/j.bbamcr.2008.10.005

Berridge, M. J., & Irvine, R. F. (1984). Inositol trisphosphate, a novel second messenger in cellular signal transduction. Nature, 312(5992), 315–321. 10.1038/312315a0

Berridge, M. J., & Irvine, R. F. (1989). Inositol phosphates and cell signalling. Nature, 341(6239), 197–205. 10.1038/341197a0

Burke, J. E., Triscott, J., Emerling, B. M., & Hammond, G. R. V. (2023). Beyond PI3Ks: Targeting phosphoinositide kinases in disease. Nature Reviews Drug Discovery, 22(5), 357–386. 10.1038/s41573-022-00582-5

de la Cruz, L., Kushmerick, C., Sullivan, J. M., Kruse, M., & Vivas, O. (2022). Hippocampal neurons maintain a large PtdIns(4)P pool that results in faster PtdIns(4,5)P2 synthesis. The Journal of General Physiology, 154(3), e202113001. 10.1085/jgp.202113001

Dickson, E. J., Falkenburger, B. H., & Hille, B. (2013). Quantitative properties and receptor reserve of the IP(3) and calcium branch of G(q)-coupled receptor signaling. The Journal of General Physiology, 141(5), 521–535. 10.1085/jgp.201210886

Falkenburger, B. H., Dickson, E. J., & Hille, B. (2013). Quantitative properties and receptor reserve of the DAG and PKC branch of G(q)-coupled receptor signaling. The Journal of General Physiology, 141(5), 537–555. 10.1085/jgp.201210887

Falkenburger, B. H., Jensen, J. B., & Hille, B. (2010). Kinetics of PIP2 metabolism and KCNQ2/3 channel regulation studied with a voltage-sensitive phosphatase in living cells. Journal of General Physiology, 135(2), 99–114. 10.1085/jgp.200910345

Grabon, A., Orłowski, A., Tripathi, A., Vuorio, J., Javanainen, M., Róg, T., Lönnfors, M., McDermott, M. I., Siebert, G., Somerharju, P., Vattulainen, I., & Bankaitis, V. A. (2017). Dynamics and energetics of the mammalian phosphatidylinositol transfer protein phospholipid exchange cycle. Journal of Biological Chemistry, 292(35), 14438–14455. 10.1074/jbc.M117.791467

Hernandez-Hernandez, G., O’Dwyer, S. C., Yang, P.-C., Matsumoto, C., Tieu, M., Fong, Z., Lewis, T. J., Santana, L. F., & Clancy, C. E. (2024). A computational model predicts sex-specific responses to calcium channel blockers in mammalian mesenteric vascular smooth muscle. eLife, 12, RP90604. 10.7554/eLife.90604

Kernik, D. C., Morotti, S., Wu, H., Garg, P., Duff, H. J., Kurokawa, J., Jalife, J., Wu, J. C., Grandi, E., & Clancy, C. E. (2019). A computational model of induced pluripotent stem-cell derived cardiomyocytes incorporating experimental variability from multiple data sources. The Journal of Physiology, 597(17), 4533–4564. 10.1113/JP277724

Kim, Y. J., Guzman-Hernandez, M.-L., Wisniewski, E., & Balla, T. (2015). Phosphatidylinositol-Phosphatidic Acid Exchange by Nir2 at ER-PM Contact Sites Maintains Phosphoinositide Signaling Competence. Developmental Cell, 33(5), 549–561. 10.1016/j.devcel.2015.04.028

Kruse, M., Vivas, O., Traynor-Kaplan, A., & Hille, B. (2016). Dynamics of Phosphoinositide-Dependent Signaling in Sympathetic Neurons. The Journal of Neuroscience, 36(4), 1386–1400. 10.1523/JNEUROSCI.3535-15.2016

Lu, J., Dong, W., Hammond, G. R., & Hong, Y. (2022). Hypoxia controls plasma membrane targeting of polarity proteins by dynamic turnover of PI4P and PI(4,5)P2. eLife, 11, e79582. 10.7554/eLife.79582

Moreno, J. D., Lewis, T. J., & Clancy, C. E. (2016). Parameterization for In-Silico Modeling of Ion Channel Interactions with Drugs. PloS One, 11(3), e0150761. 10.1371/journal.pone.0150761

Morleo, M., Venditti, R., Theodorou, E., Briere, L. C., Rosello, M., Tirozzi, A., Tammaro, R., Al-Badri, N., High, F. A., Shi, J., Acosta, M. T., Adam, M., Adams, D. R., Alvarez, R. L., Alvey, J., Amendola, L., Andrews, A., Ashley, E. A., Bacino, C. A., … Franco, B. (2023). De novo missense variants in phosphatidylinositol kinase PIP5KIγ underlie a neurodevelopmental syndrome associated with altered phosphoinositide signaling. The American Journal of Human Genetics, 110(8), 1377–1393. 10.1016/j.ajhg.2023.06.012

Nakatsu, F., Baskin, J. M., Chung, J., Tanner, L. B., Shui, G., Lee, S. Y., Pirruccello, M., Hao, M., Ingolia, N. T., Wenk, M. R., & De Camilli, P. (2012). PtdIns4P synthesis by PI4KIIIα at the plasma membrane and its impact on plasma membrane identity. Journal of Cell Biology, 199(6), 1003–1016. 10.1083/jcb.201206095

Narkis, G., Ofir, R., Landau, D., Manor, E., Volokita, M., Hershkowitz, R., Elbedour, K., & Birk, O. S. (2007). Lethal Contractural Syndrome Type 3 (LCCS3) Is Caused by a Mutation in *PIP5K1C*, Which Encodes PIPKIγ of the Phophatidylinsitol Pathway. The American Journal of Human Genetics, 81(3), 530–539. 10.1086/520771

Pemberton, J. G., Kim, Y. J., & Balla, T. (2020). Integrated regulation of the phosphatidylinositol cycle and phosphoinositide-driven lipid transport at ER-PM contact sites. Traffic, 21(2), 200–219. 10.1111/tra.12709

Rhana, P., Matsumoto, C., Fong, Z., Costa, A. D., Del Villar, S. G., Dixon, R. E., & Santana, L. F. (2024). Fueling the heartbeat: Dynamic regulation of intracellular ATP during excitation-contraction coupling in ventricular myocytes. Proceedings of the National Academy of Sciences of the United States of America, 121(25), e2318535121. 10.1073/pnas.2318535121

Salter, C. G., Cai, Y., Lo, B., Helman, G., Taylor, H., McCartney, A., Leslie, J. S., Accogli, A., Zara, F., Traverso, M., Fasham, J., Lees, J. A., Ferla, M. P., Chioza, B. A., Wenger, O., Scott, E., Cross, H. E., Crawford, J., Warshawsky, I., … Baple, E. L. (2021). Biallelic PI4KA variants cause neurological, intestinal and immunological disease. Brain, 144(12), 3597–3610. 10.1093/brain/awab313

Suresh, S., Shaw, A. L., Pemberton, J. G., Scott, M. K., Harris, N. J., Parson, M. A. H., Jenkins, M. L., Rohilla, P., Alvarez-Prats, A., Balla, T., Yip, C. K., & Burke, J. E. (2024). Molecular basis for plasma membrane recruitment of PI4KA by EFR3. Science Advances, 10(51), eadp6660. 10.1126/sciadv.adp6660

Verdura, E., Rodríguez-Palmero, A., Vélez-Santamaria, V., Planas-Serra, L., de la Calle, I., Raspall-Chaure, M., Roubertie, A., Benkirane, M., Saettini, F., Pavinato, L., Mandrile, G., O’Leary, M., O’Heir, E., Barredo, E., Chacón, A., Michaud, V., Goizet, C., Ruiz, M., Schlüter, A., … Pujol, A. (2021). Biallelic PI4KA variants cause a novel neurodevelopmental syndrome with hypomyelinating leukodystrophy. Brain, 144(9), 2659–2669. 10.1093/brain/awab124

Viaud, J., Mansour, R., Antkowiak, A., Mujalli, A., Valet, C., Chicanne, G., Xuereb, J.-M., Terrisse, A.-D., Séverin, S., Gratacap, M.-P., Gaits-Iacovoni, F., & Payrastre, B. (2016). Phosphoinositides: Important lipids in the coordination of cell dynamics. Biochimie, 125, 250–258. 10.1016/j.biochi.2015.09.005

Wang, D. G., Paddock, M. N., Lundquist, M. R., Sun, J. Y., Mashadova, O., Amadiume, S., Bumpus, T. W., Hodakoski, C., Hopkins, B. D., Fine, M., Hill, A., Yang, T. J., Baskin, J. M., Dow, L. E., & Cantley, L. C. (2019). PIP4Ks Suppress Insulin Signaling through a Catalytic-Independent Mechanism. Cell Reports, 27(7), 1991–2001.e5. 10.1016/j.celrep.2019.04.070

Willars, G. B., Nahorski, S. R., & Challiss, R. A. J. (1998). Differential Regulation of Muscarinic Acetylcholine Receptor-sensitive Polyphosphoinositide Pools and Consequences for Signaling in Human Neuroblastoma Cells*. Journal of Biological Chemistry, 273(9), 5037–5046. 10.1074/jbc.273.9.5037

Wills, R. C., Doyle, C. P., Zewe, J. P., Pacheco, J., Hansen, S. D., & Hammond, G. R. V. (2023). A novel homeostatic mechanism tunes PI(4,5)P2-dependent signaling at the plasma membrane. Journal of Cell Science, 136(16), jcs261494. 10.1242/jcs.261494

Xu, C., Watras, J., & Loew, L. M. (2003). Kinetic analysis of receptor-activated phosphoinositide turnover. The Journal of Cell Biology, 161(4), 779–791. 10.1083/jcb.200301070

Zandl-Lang, M., Plecko, B., & Köfeler, H. (2023). Lipidomics—Paving the Road towards Better Insight and Precision Medicine in Rare Metabolic Diseases. International Journal of Molecular Sciences, 24(2), Article 2. 10.3390/ijms24021709

